# Allosteric receptor modulation of FFA2R turns natural agonists into potent activators of the superoxide generating neutrophil NADPH-oxidase

**DOI:** 10.1101/284935

**Authors:** Jonas Mårtensson, André Holdfeldt, Martina Sundqvist, Michael Gabl, Lena Björkman, Huamei Forsman, Claes Dahlgren

## Abstract

Acetate, agonist for the free fatty acid receptor 2 (FFA2R/GPR43), triggers an increase in the cytosolic concentration of free Ca^2+^ in neutrophils without any assembly of the superoxide generating NADPH-oxidase. We show that the phenylacetamide compound 58 (Cmp58; (S)-2-(4-chlorophenyl)-3,3-dimethyl-N-(5-phenylthiazol-2-yl)butanamide, lacking a direct activating effect on neutrophils, acts as a positive allosteric FFA2R modulator that turns acetate into a potent activating agonist that triggers an assembly of the NADPH-oxidase. The NADPH-oxidase activity could be further increased in neutrophils treated with the pro-inflammatory cytokine TNF. Many neutrophil chemoattractant receptors are stored in secretory organelles but no FFA2R mobilization was induced in neutrophils treated with TNF. The receptor selectivity was demonstrated through the inhibition of the neutrophil response induced by the combined action of acetate and Cmp58 by the FFA2R antagonist CATPB. Allosteric modulators that positively co-operate with natural FFA2R agonists and prime neutrophils in their response to such agonists, may serve as good tools for further unraveling the physiological functions of the FFA2R and its involvement in various diseases. In this study, allosteric modulation of FFA2R is introduced as a novel receptor selective mechanism to prime neutrophils to produce increased amounts of reactive oxygen species.

## Introduction

An inflammatory response is normally initiated, maintained, and sometimes modulated by microbial or host derived pathogen- or danger-associated molecular patterns (PAMPs and DAMPs) (1–3). It is clear that these DAMPs/PAMPs together participate in a multistep hierarchy of directional cues that regulate neutrophil function at inflammatory sites (4, 5). The importance of microbial products in regulating homeostasis and inflammatory responses in the gut is well known, and the number/amount of more or less well characterized and possibly also unknown metabolites released from growing bacteria that should be dealt with is very large. Peptides containing a formylated methionine (fMet) in their N-terminus constitute one such group of microbial derived “molecular pattern” which are recognized by cells of our innate immune system with the rational being that bacterial protein synthesis starts with an fMet (6). Short chain fatty acids (SCFA), produced through microbial fermentation of fiber-rich diets, are also of specific interest in relation to inflammatory diseases associated with the gut microbiota (7, 8). Products of bacterial metabolism such as formyl peptides and short chain fatty acids (SCFAs) exert their functions in relation to cells in our innate immune system through their binding to pattern recognition receptors (PRRs); formylated peptides being recognized by the formyl peptide receptors (FPRs; FPR1 and FPR2) (9–11)) whereas the SCFAs are recognized by two until fairly recently orphan receptors, FFA2R and FFA3R (earlier known as GPR43 and GPR41, respectively; (12, 13)). The FPRs as well as the FFARs belong to the family of G-protein coupled receptors (GPCRs (14)).

The SCFAs produced by bacteria can reach very high concentrations in the gut (15, 16), and they are subsequently transported over the gut epithelium to be released in the bloodstream where they may reach millimolar concentration levels. The SCFA receptor FFA3R (GPR41) has been shown not to be expressed in neutrophils whereas FFA2R is highly expressed in these cells (17), and published data suggest that the receptor has important roles in orchestration of compound 58 (Cmp58; (S)-2-(4-chlorophenyl)-3,3-dimethyl-N-(5-phenylthiazol-2-yl)butanamide) is one of them and it was recently classified as an allosteric FFA2R agonist/modulator, when shown to change the efficacy of the response induced by acetate as measured in a forskolin-induced cAMP assay system (22). In this study, we have investigated the effects of this allosteric FFA2R agonist/modulator on neutrophils. Our data show that Cmp58 functions as a positive allosteric modulator in these cells by turning the natural FFA2R agonist acetate into a potent activator of the neutrophil NADPH-oxidase. A novel receptor selective neutrophil priming mechanism is, thus, disclosed.

## Material and Methods

### Chemicals

The peptides fMLF and WKYMVM were synthesized and HPLC-purified by TAG Copenhagen A/S (Copenhagen, Denmark). Isoluminol, TNF-*α*, the FFA3R agonist AR420626, PMA (phorbol 12-myristate 13-acetate), propionic acid, and pertussis toxin (PTX) were purchased from Sigma (Sigma Chemical Co., St. Louis, MO, USA). Cyclosporin H was a kind gift provided by Novartis Pharma (Basel, Switzerland). Dextran and Ficoll-Paque were obtained from GE-Healthcare Bio-Science (Uppsala, Sweden). Fura 2-AM was from Molecular Probes/Thermo Fisher Scientific (Waltham, MA, USA), and horseradish peroxidase (HRP) was obtained from Boehringer Mannheim (Mannheim, Germany). The phenylacetamide compound (S)-2-(4-chlorophenyl)-3,3-dimethyl-N-(5-phenylthiazol-2-yl)butanamide (PA;Cmp58) was obtained from Calbiochem-Merck Millipore (Billerica, USA; (22)) and 4-chloro-α-(1-methylethyl)-N-2-thiazolyl-benzeneacetamide (4-CMTB) was from TOCRIS (Bristol, UK).

Peptides were dissolved in DMSO and stored at -80°C until use. Subsequent dilutions of all peptides and other reagents were made in Krebs-Ringer Glucose phosphate buffer (KRG; 120 mM NaCl, 4.9 mM KCl, 1.7 mM KH_2_PO_4_, 8.3 mM NaH_2_PO_4_, 1.2 mM MgSO_4_, 10 mM glucose, and 1 mM CaCl_2_ in dH_2_O, pH 7.3). Ammonium acetate (NH_4_OAc) was from Merck (Germany), and the Gαq inhibitor YM-254890 was purchased from Wako Chemicals (Neuss, Germany). The phycoerythrine (PE) conjugated antibodies against CD11b were from Becton Dickinson (New Jersey, USA) and against FFA2R/GPR43 were from Bios (Woburn, USA). The FFA2R agonist Cmp1 (3-benzyl-4-(cyclopropyl-(4-(2,5-dichlorophenyl)thiazol-2-yl)amino)-4-oxobutanoic acid and the antagonist CATPB ((S)-3-(2-(3-chlorophenyl)acetamido)-4-(4-(trifluoromethyl)-phenyl) butanoic acid) synthesized as described previously (20, 31, 32), were generous gifts from Trond Ulven (Odense university, Denmark).

### Isolation of neutrophils from human blood

Neutrophils were isolated from buffy coats from healthy blood donors by dextran sedimentation and Ficoll-Paque gradient centrifugation as described by Bøyum (33). Remaining erythrocytes were removed by hypotonic lysis and the neutrophils were washed and resuspended in KRG and stored on ice until use.

### Measuring NADPH-oxidase activity

Isoluminol-enhanced chemiluminescence (CL) technique was used to measure superoxide production, the precursor of production of reactive oxygen species (ROS), by the NADPH-oxidase activity as described (34, 35). In short, the CL measurements were performed in a six-channel Biolumat LB 9505 (Berthold Co., Wildbad, Germany), using disposable 4-ml polypropylene tubes and a 900 *μ*l reaction mixture containing 10^5^ neutrophils, isoluminol (2×10^−5^ M) and HRP (4 Units/mL). The tubes were equilibrated for 5 min at 37°C, before addition of agonist (100 *μ*l) and the light emission was recorded continuously over time. In experiments where the effects of receptor specific antagonists were determined, these were added to the reaction mixture 1-5 min before stimulation with control neutrophils incubated under the same condition but in the absence of antagonist run in parallel for comparison. To determine the involvement of G*α*i, neutrophils (1×10^6^/ml) in KRG were incubated without and with PTX (500 ng/ml final concentration) at 37°C for different periods of time and the neutrophil superoxide production was then determined.

### Calcium mobilization

Neutrophils at a density of 1–3×10^6^ cells/ml were washed with Ca^2+^-free KRG and centrifuged at 220x*g*. The cell pellets were re-suspended at a density of 2×10^7^ cells/ml in KRG containing 0.1% BSA, and loaded with 2 *μ*M FURA 2-AM for 30 minutes at room temperature. The cells were then washed once with KRG and resuspended in the same buffer at a density of 2×10^7^cells/ml. Calcium measurements were carried out in a Perkin Elmer fluorescence spectrophotometer (LC50), with excitation wavelengths of 340 nm and 380 nm, an emission wavelength of 509 nm, and slit widths of 5 nm and 10 nm, respectively. The transient rise in intracellular calcium is presented as the ratio of fluorescence intensities (340 nm / 380 nm) detected.

### Cell-surface receptor expression

The level of surface expression of the integrin CR3 (CD11b) and FFA2R was determined by the use of PE-conjugated antibodies against CD11b and FFA2R, respectively. Briefly, neutrophils (5 *×* 10^6^/mL) in KRG were incubated at 37°C in the absence or presence of TNF-*α* (10 ng/mL; 20 min) before addition of antibodies and incubation on ice for 30 min in darkness. Thereafter, cells were washed, to remove excess unbound antibodies, resuspended in KRG and kept on ice in darkness until the amount of bound fluorescence was determined by flow cytometry using an Accuri C6 flow cytometer (Becton Dickinson Sparks, MD, USA).

## Results

### Allosteric modulation of FFA2R lowers the concentration of acetate required to induce a transient rise in the concentration of intracellular Ca^2+^ in neutrophils

The receptor for acetate, FFA2R, belongs to the family of G-protein coupled receptors (GPCRs). A very early signaling route in neutrophils, generated downstream of many agonist activated GPCRs, is an increase in the cytosolic concentration of free Ca^2+^ ([Ca^2+^]_i_) (36), triggered through the Gi*β*/*γ*-complex or the Gq*α*-subunit mediated PLC activation and cleavage of PIP_2_ (phosphatidyl-inositol-4,5-bis-phosphate) giving rise to inositol-1,4,5-tris-phosphate (IP_3_) and a transient Ca^2+^ rise. Accordingly, the FFA2R agonist acetate induced a concentration-dependent transient increase in [Ca^2+^]_i__i_ when interacting with neutrophils (Fig 1 upper panel and (23)). No such direct activating effect was induced by the phenylacetamide compound58 (Cmp58) or another earlier described allosteric FFA2R agonist, 4-CMTB (29), investigated in concentrations up to 5*μ*M (shown for Cmp58 in Fig 1 middle panel including also the response induced by the FPR1 agonist fMLF for comparison). These compounds specifically recognizes an allosteric FFA2R site and modulates receptor function (22, 29). The suggested positively modulatory effect mediated by Cmp58 was clearly evident by comparing the neutrophil response when stimulated with acetate alone (Fig 1 upper panel) and in the presence of Cmp58 (Fig 1 lower panel). At concentration where no rise in [Ca^2+^]_i__i_ could be detected with acetate alone, neutrophils responded well when first incubated with Cmp58 (Fig 1 lower panel). In summary, Cmp58 lowered the thresh-hold for the acetate-induced response as illustrated by the fact that concentrations of the natural FFA2R agonist acetate that were non-activating in naïve neutrophils induced a transient rise in the cytosolic concentration of free Ca^2+^ in cells pre-treated with Cmp58 (Fig 1).

**Figure 1.**
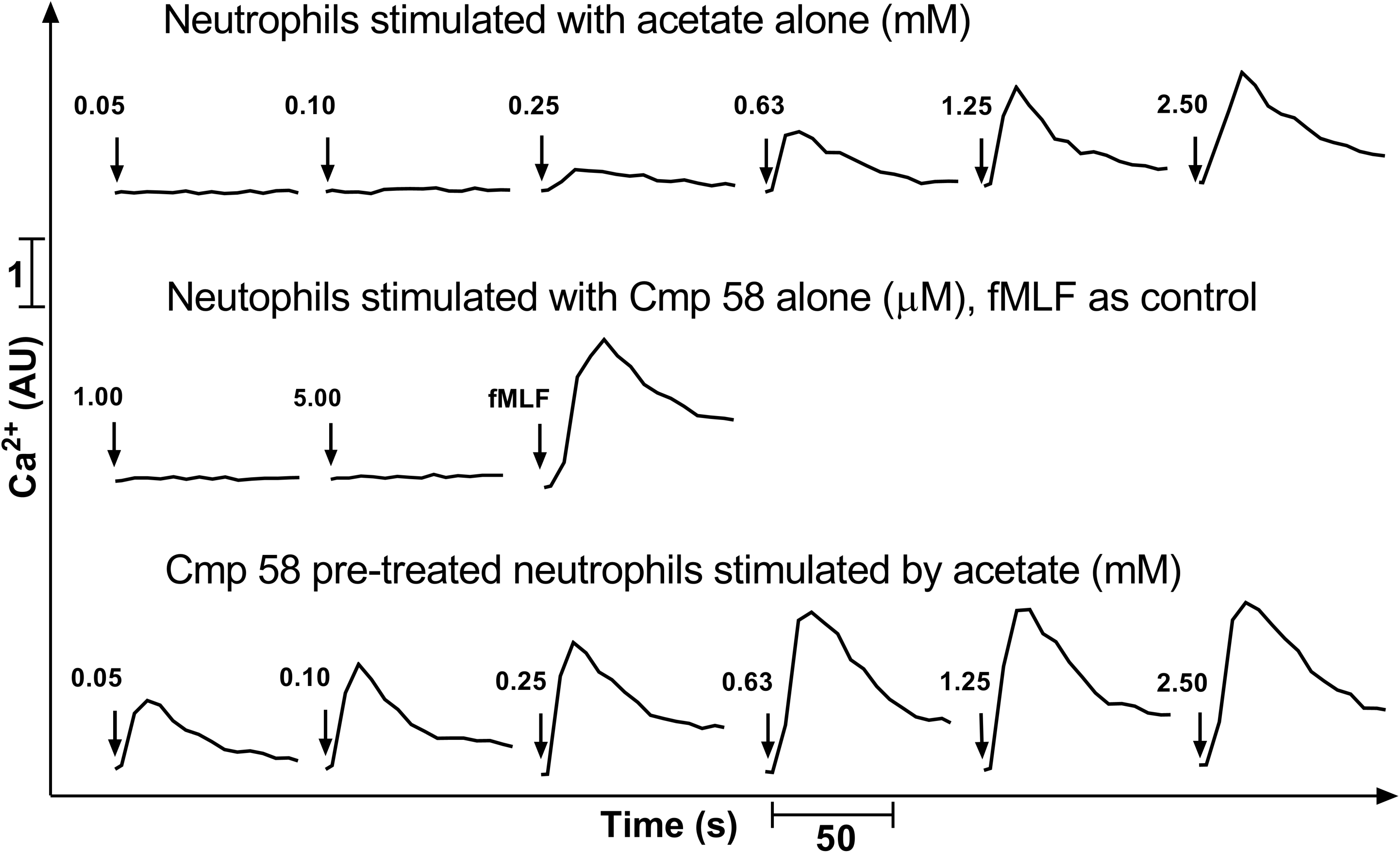
The allosteric FFA2R agonist Cmp58 modulates the transient rise in intracellular Ca^2+^ induced by acetate in neutrophils. Human neutrophils loaded with Fura-2 were stimulated with acetate (upper panel; different concentrations) or Cmp58 and acetate added in sequence (lower panel; first Cmp58 1*μ*M, and thereafter different concentrations of acetate). No neutrophil response was induced by Cmp58 alone (middle panel) and for comparison, the neutrophil response induced by the FPR1 agonist (fMLF; 5nM) is also shown. The time point for addition of an agonist is indicated by an arrow. Abscissa, time of study (sec); Ordinate, Increase in intracellular [Ca^2+^]_i_ given by the ratio between Fura-2 fluorescence at 340 and 380 nm.

### Allosteric modulation of FFA2R turns acetate to a highly efficient activator of the neutrophil NADPH-oxidase

In addition to the triggering of the PLC-PIP_2_-IP_3_ signaling route that gives rise to a transient increase in [Ca^2+^]_i_, many GPCR agonists also activate the NADPH-oxidase, an electron transporting enzyme system in neutrophils that generates superoxide anions (O_2_^−^; (37)). In agreement with the inability to mobilize Ca^2+^ (Fig 1 middle panel), Cmp58 had no direct neutrophil activating effects on the oxidase system as no superoxide release was induced in neutrophils allowed to interact with the FFA2R modulator (Fig 2). Also the natural FFA2R agonist acetate was shown to be a poor activator of the neutrophil NADPH-oxidase (Fig 2; see also (23)), as compared to the responses induced by the FPR agonists fMLF (FPR1 agonist) and WKYMVM (FPR2 agonist; Fig 2). However, when combined with Cmp58, the NADPH-oxidase activity induced by acetate was largely augmented (Fig 3). The increased production/release of superoxide was evident when Cmp58 was added concomitantly with acetate (Fig 3A), but even more so when neutrophils were pre-treated for five min with Cmp58 before stimulation with acetate (Fig 3C). The amplifying effect with the two FFA2R ligands working together was evident even when the order of addition was reversed (first acetate and then Cmp58) but the amount of superoxide released was lower than when the order was first Cmp58 and then acetate (Fig 3B). In addition, a primed response to acetate was achieved also when 4-CMTB replaced Cmp58 as the allosteric amplifier, but the priming effect was substantially lower than that induced by Cmp58 (Fig 3C inset).

**Figure 2.**
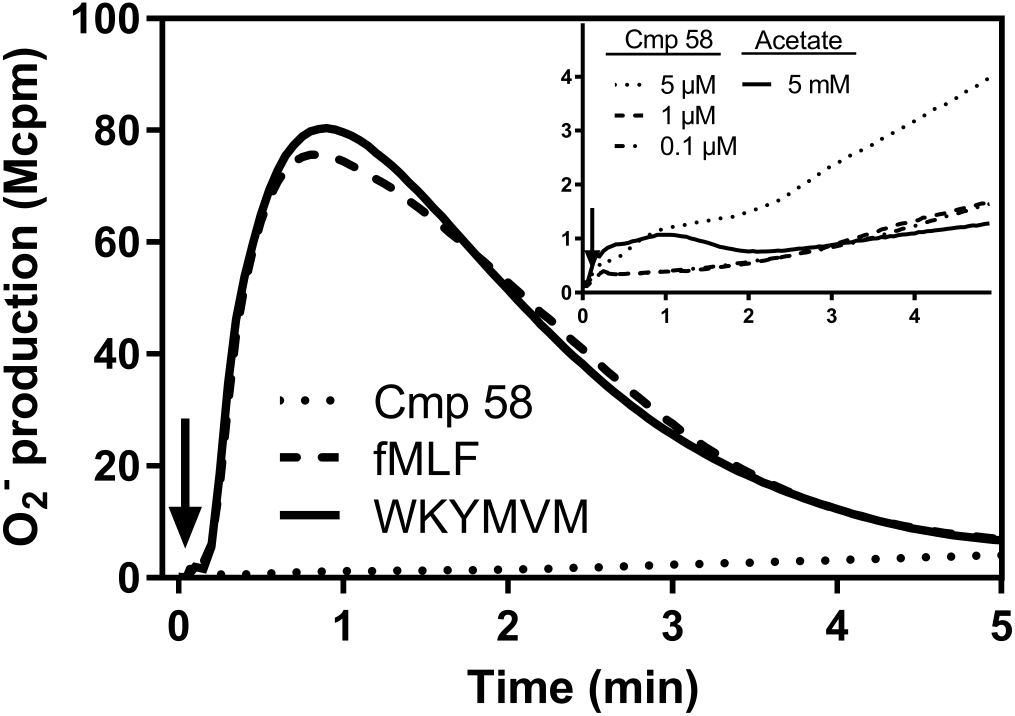
The allosteric FFA2R ligand Cmp58 has no direct effect on the neutrophil NADPH-oxidase activity. Human neutrophils were activated with Cmp58 alone (5 *μ*M, dotted line) and the release of superoxide anions was recorded. One representative experiment out of >5 is shown, and the responses induced by the FPR agonists fMFL (100 nM; FPR1 agonist, solid line) and WKYMVM (100 nM; FPR2 agonist, dashed line) are included for comparison. Superoxide production was recorded continuously and the time point for addition of the agonists is indicated by an arrow. Abscissa, Time of study (min); Ordinate, Superoxide production (arbitrary units; Mcpm). Inset: The neutrophil response induced by different concentrations of Cmp58 and acetate alone (5mM) are shown and the time point for addition of Cmp58/acetate is indicated by an arrow. Abscissa, Time of study (min); Ordinate, Superoxide production (arbitrary units; Mcpm)

**Figure 3.**
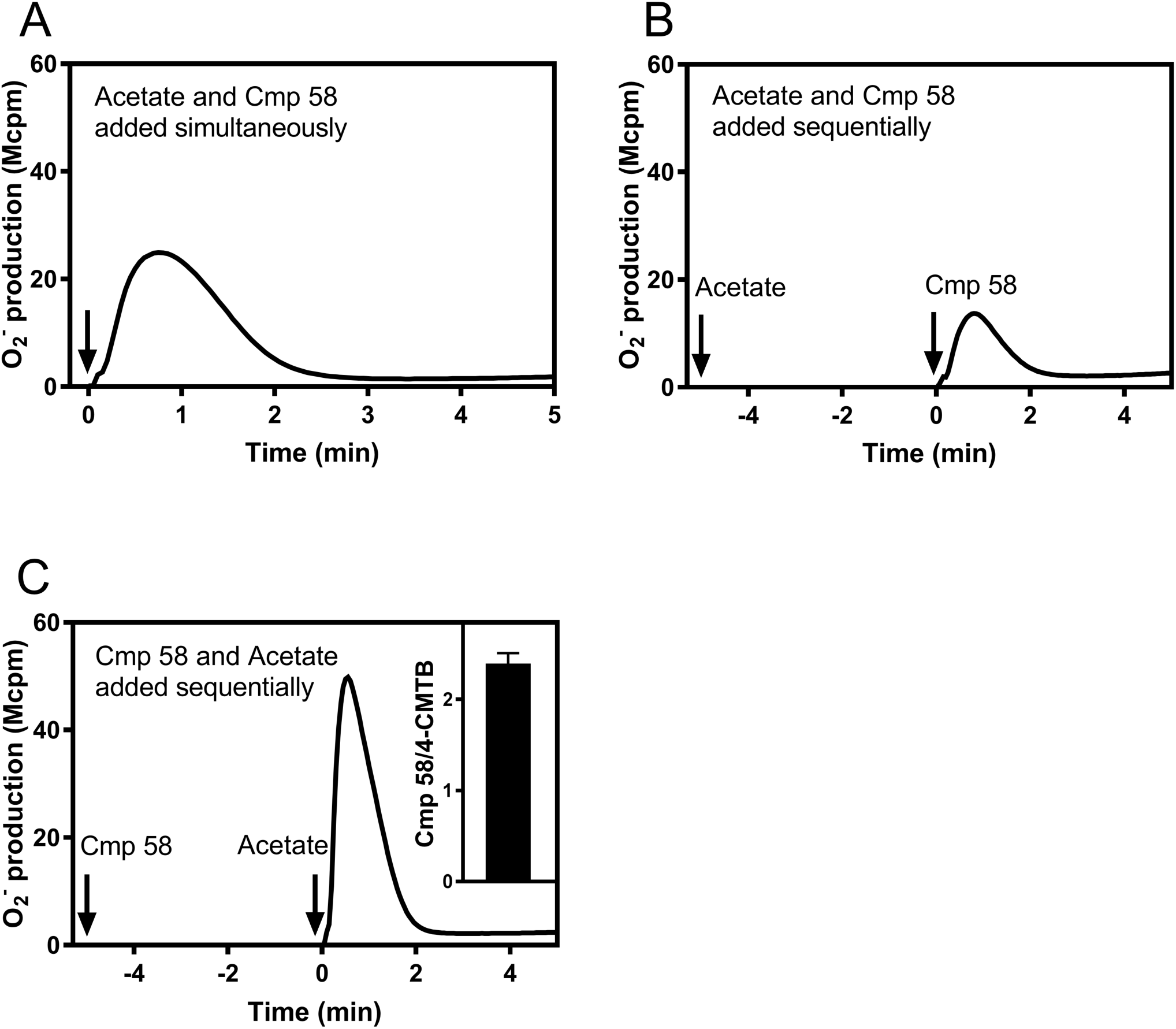
Neutrophil activation of the NADPH-oxidase by Cmp58 and acetate when added together or in sequence. Human neutrophils were activated with Cmp58 (1*μ*M) and acetate (1mM) and the two FFA2R ligands were added simultaneously (A) or in sequence (B and C). The time points for addition of the molecules are indicated by arrows and superoxide production was recorded continuously. Abscissa, Time of study (min); Ordinate, Superoxide production (arbitrary units; Mcpm). Inset in C: The bar graph shows a comparison of the response induced by acetate (1mM) in neutrophils treated for 5 min with Cmp58 (1*μ*M) or 4-CMTB (1*μ*M). The peak responses were determined and the results are given as the ratio between the activities in the presence of 4-CMTB and Cmp58 (mean±SD; n=3).

The increase of neutrophil superoxide release was most pronounced with Cmp58 and when the cells were pre-incubated with the modulator prior to the addition of acetate, and in order to characterize the response, we used the most potent modulator (Cmp58) and partly “clamped” the system by keeping the concentration of the modulator constant (1*μ*M) and the priming (incubation) time fixed to five min. Using this setup, it is clear that superoxide release from neutrophils activated with acetate was dependent on the concentration of the orthosteric FFA2R agonist with an EC_50_ value of ≈100*μ*M and a full response at around 500*μ*M (Fig 4A). When using an acetate concentration giving a maximum response (1mM) neutrophil superoxide release was dependent on the concentration of Cmp58 with an EC_50_ value of ≈250nM, reaching a full response with a concentration of around 1*μ*M (Fig 4B). Regarding the priming time needed to obtain a good modulating effect, time titration experiments showed that a less than two minutes pre-incubation with Cmp58 was sufficient to achieve a full neutrophil response upon addition of acetate (Fig 4C).

**Figure 4.**
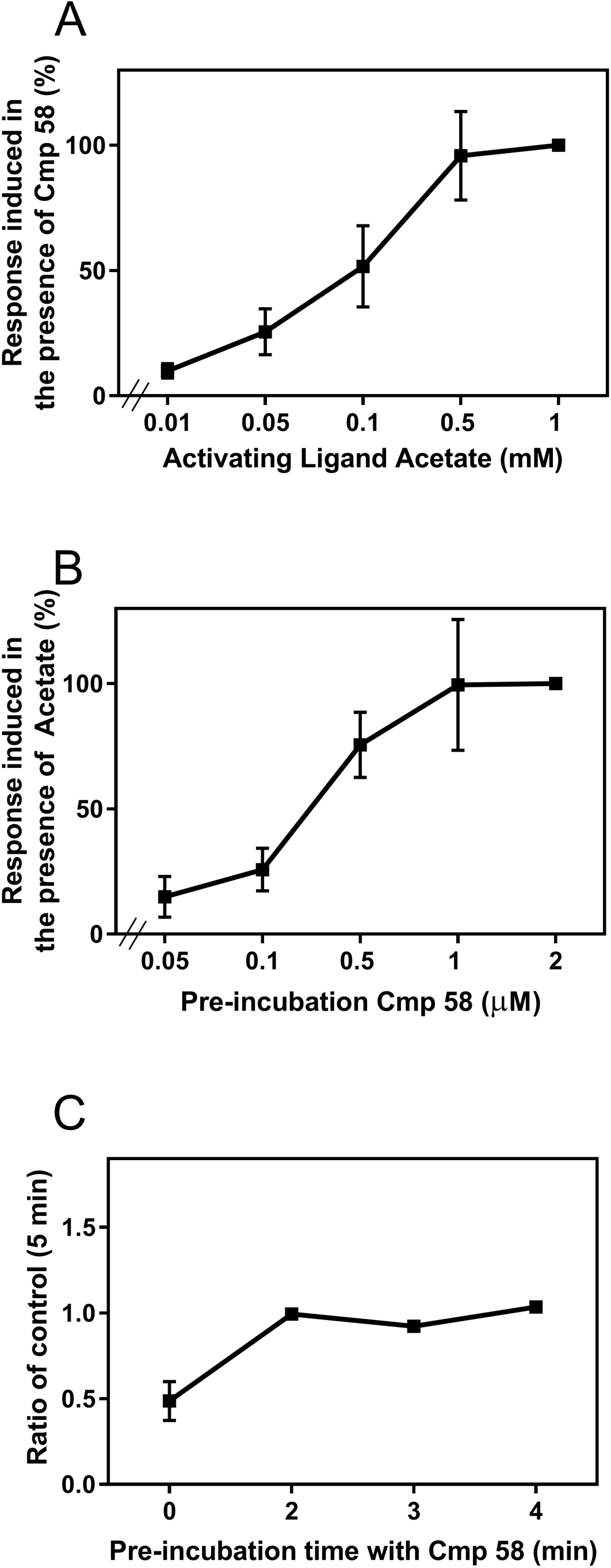
Dose and time titration of NADPH-oxidase activity induced in neutrophils stimulated with acetate after preincubation with Cmp58. Human neutrophils were incubated without and with Cmp58 (1*μ*M) for 5 min at 37°C followed by an activation with acetate at different concentrations. Superoxide production was recorded continuously and the peak values of the responses were determined and compared. Abscissa, Concentration of acetate (mM); Ordinate, Superoxide production expressed in percent of the activity induced by 1mM acetate (mean±SD; n=3). A. Human neutrophils were incubated without and with different concentrations of Cmp58 as indicated for 5 min at 37°C and then activated by and acetate (1mM). Superoxide production was recorded continuously and the peak values of the responses were determined and compared. Abscissa, Concentration of Cmp58 (*μ*M); Ordinate, Superoxide production given in percent of the activity induced by acetate with neutrophils preincubated with 1*μ*M Cmp58 (mean±SD; n=3). B. Human neutrophils were incubated with Cmp58 (1*μ*M) for up to 5 min at 37°C and the cells were then stimulated with acetate (1mM). Superoxide production was recorded continuously and the peak values of the responses were determined and compared to the value obtained with neutrophils pre-incubated for 5 min with Cmp58. Abscissa, pre-incubation time (min); Ordinate, Superoxide production given as ratios with the response induced in neutrophils preincubated with Cmp58 for 5 min (mean±SD; n=3).

In quantitative terms the neutrophil superoxide production induced by 1mM acetate in neutrophils pretreated for five minutes with 1*μ*M Cmp58 was of the same magnitude as that induced by the commonly used FPR agonists fMLF and WKYMVM (100nM; Fig 5A). It was also clear that the response induced by the fairly high concentration of FPR agonists were not affected by pre-incubation with Cmp58 (Fig 5B).

**Figure 5.**
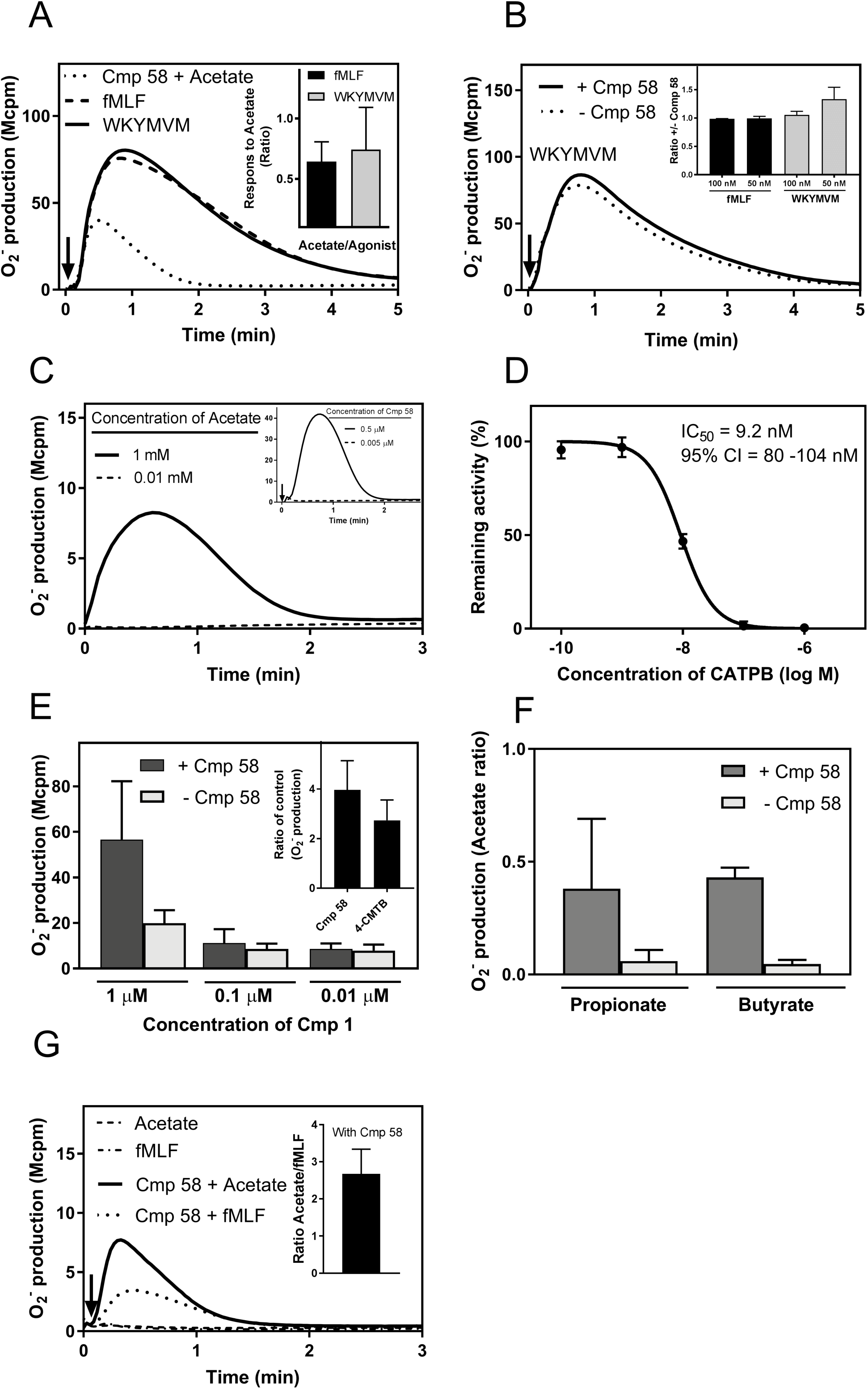
Basic characteristics of the augmented NADPH-oxidase activity mediated by Cmp58. A. Human neutrophils were incubated with Cmp58 (1*μ*M) for 5 min at 37°C and then activated by acetate (1mM; dotted line). Superoxide production was recorded continuously and the response was compared to the activities induced by fMLF (100nM; FPR1 agonist; broken line) and WKYMVM (100nM; FPR2 agonist; solid line). The time point for addition of the agonists is indicated by an arrow. Abscissa, Time of study (min); Ordinate, Superoxide production (in arbitrary units; Mcpm). Inset: Human neutrophils were incubated with Cmp58 (1*μ*M) for 5 min at 37°C and then activated by acetate (1mM). Superoxide production was recorded continuously and the response was compared to the activities induced by fMLF (100nM; FPR1 agonist) and WKYMVM (100nM; FPR2 agonist). The peak values of the responses were determined, compared, and are presented as the ratios WKYMVM/Cmp58+acetate (black) and fMLF/Cmp58+acetate (grey), respectively (mean±SD; n=3). B. Human neutrophils were incubated without (dotted line) or with (solid line) Cmp58 (1*μ*M) for 5 min at 37°C and then activated with WKYMVM (100nM). Superoxide production was recorded continuously and the time point for addition of the agonists is indicated by an arrow. Abscissa, Time of study (min); Ordinate, Superoxide production (in arbitrary units; Mcpm). Inset: Human neutrophils were incubated without or with Cmp58 (1*μ*M) for 5 min at 37°C and then activated by and WKYMVM (100nM or 50nM) or fMLF (100nM or 50nM). Superoxide production was recorded continuously, the peak activities were determined and ratios with/without Cmp58 were calculated for 50 and 100nM of the FPR agonists, respectively (ratios fMLF+/-Cmp58, black and WKYMVM+/-Cmp58, grey; mean±SD, n=3). C. Human neutrophils were incubated with acetate (1mM) for 5 min and the cells were then diluted 1/200 in measurement vials containing Cmp58 (1*μ*M) without (dashed line) or with (solid line) acetate (1mM) and production of superoxide was recorded directly. One representative experiment out of 3 is shown. Abscissa, Time of study (min); Ordinate, Superoxide production (arbitrary units; (Mcpm). Inset: Human neutrophils were incubated with Cmp58 (0.5*μ*M) for 10 min at 37°C and the cells were then diluted 1/100 in measurement vial without (dashed line) or with Cmp58 (0.5*μ*M, solid line) and the cells (in 0.5 and 0.005*μ*M Cmp58) were activated by acetate 30 seconds after the samples were diluted in new buffer. Superoxide production was recorded continuously and the time point for addition of acetate is indicated by an arrow and one representative experiment out of more than 3 is shown. Abscissa, Time of study (min); Ordinate, Superoxide production (arbitrary units; (Mcpm). D. Human neutrophils were incubated with the FFA2R antagonist CATPB at different concentrations as indicated and the activated by the combination of Cmp58 (1*μ*M; 5 min together with CATPB) and acetate (1mM). The neutrophil response was measured as superoxide production that was recorded continuously. The peak activities were determined and the results expressed as percent of the activity induced by Cmp58/acetate in the absence of the FFA2R antagonist. Abscisssa, Concentration of CATPB (log M); Ordinate, remaining activity (in percent of the activity induced without any antagonist; mean±SD, n=3). E. Human neutrophils were incubated without (dark grey bars) or with Cmp 58 (1*μ*M; light grey bars) for 5 minutes and then activated with the FFA2R agonist Cmp1 at different concentrations (*μ*M) as indicated. Superoxide production was recorded continuously and the peak activities were determined. Abscissa, Concentration of Cmp1 (*μ*M); Ordinate, Superoxide production (peak value in arbitrary units; (Mcpm; mean±SD, n=3). Inset: Human neutrophils were incubated without or with Cmp 58 (1*μ*M) and 4-CMTB (5*μ*M), respectively, for 5 minutes and then activated with the FFA2R agonist Cmp1 (1*μ*M). Superoxide production was recorded continuously and the peak activities were determined and the results are given as fold increase in the presence of the modulators compared to the activity induced by Cmp1 alone. (mean±SD, n=3). F. Human neutrophils were incubated with Cmp58 (1*μ*M) for 5 min at 37°C and then activated by acetate, propionate or butyrate (1mM). Superoxide production was recorded continuously and the responses were compared. The responses induced by propionate and butyrate in the absence of Cmp58 are also shown. The peak values of the responses were determined, and given as the ratios acetate/propionate and acetate/butyrate, respectively (mean±SD; n=3). G. Human neutrophils were incubated with or without Cmp58 (1*μ*M) for 5 min at 37°C and then activated by and acetate (1mM) or fMLF (1nM). The time point for addition of the agonists is indicated by an arrow. Abscissa, Time of study (min); Ordinate, Superoxide production (in arbitrary units; Mcpm). Inset: Human neutrophils were incubated with Cmp58 (1*μ*M) for 5 min at 37°C and then activated by acetate (1mM) or fMLF (1nM). Superoxide production was recorded continuously and the responses were compared. The peak values of the responses were determined, and is shown as the ratio Cmp58+acetate/Cmp58+fMLF (mean±SEM; n=6).

In order to determine if the presence of high concentration of acetate was needed to activate the oxidase we performed a “wash out” experiment. In brief, neutrophils (10^7^/ml) were pre-incubated with acetate (1mM) and the cell/acetate mixture was then diluted in pre-warmed measuring vials containing Cmp58 (1 *μ*M), without or combined with acetate (1mM), giving a final concentration of acetate in the cell-samples of 1mM and 0.01 mM, respectively. The NADPH-oxidase activity was determined directly and, whereas the NADPH-oxidase was activated in neutrophils exposed to 1mM acetate during the whole procedure, no response was induced when the concentration of acetate was rapidly reduced to 0.01mM; Fig 5C). Our data thus show that the effect of acetate is rapidly (or rather immediately) lost upon dilution, suggesting that the dissociation of bound agonist (off-rate) is very rapid.

### The primed acetate response was inhibited by the FFA2R antagonist CATPB and the allosteric modulator primes also the neutrophil response induced by other FFA2R agonists

Next we used the FFA2R antagonist CATPB (20), earlier shown to inhibit FFA2R in neutrophils (23), to further determine the receptor involved in the oxidase activity induced by acetate in cells primed with Cmp58. CATPB potently inhibited the response, shown by the fact that the neutrophil activity induced by 1mM acetate was fully inhibited by 0.1*μ*M CATPB (Fig 5D). The inhibitory effects were the same when CATPB was added prior to Cmp58 and also three minutes after Cmp58 (data not shown).

Small synthetic molecule ligands with improved potency and selectivity for FFA2R have recently become available and we have previously shown that one such specific receptor agonist (Cmp1) triggers an assembly of the NADPH-oxidase and release of superoxide anions in neutrophils (23). Using Cmp58, we now show that apart from the natural agonist acetate, the neutrophil response induced by Cmp1 was also enhanced by Cmp58 (Fig 5E), and this feature was true also for two other natural FFA2R agonists propionate and butyrate were used (Fig 5F). Further, the same pattern was observed when Cmp58 was replaced by 4-CMTB as shown for Cmp1 induced superoxide release, but the modulating effect of 4-CMTB was lower than that of Cmp58 (Inset Fig 5E). Very low levels of neutrophil NADPH-oxidase activity were induced by propionate/butyrate alone (investigated with concentrations up to 5mM) and the Cmp58 primed responses induced by the longer fatty acids were lower than that induced by acetate (Fig 5F). The receptor selectivity of the allosteric effect of Cmp58 was investigated and no neutrophil NADPH-oxidase activity was induced in the absence or presence of Cmp58 by the FFA3R selective agonist AR420626 (determined with concentrations up to 5*μ*M; data not shown). The receptor selectivity was, however, not absolute as indicated by the results obtained with low concentrations of the FPR1 agonist fMLF. A neutrophil NADPH-oxidase activity was induced in the presence of Cmp58 when neutrophils were activated by a concentration of the FPR1 that was non-activating in the absence of the modulator. The Cmp58 mediated augmentation of the fMLF response was not as pronounced as that of the acetate response (Fig 5G).

Our data show that the pre-incubation time with Cmp58 required to obtain a full neutrophil response upon addition of acetate is very short (Fig 5C). In order to determine if the effect of Cmp58 is reversible, neutrophils (10^7^/ml) were pre-incubated with Cmp58 and the cell/compound mixture was then diluted in pre-warmed measuring vials with or without Cmp58 (0.5 *μ*M), i.e., the final concentration of Cmp58 in these cell-samples were 0.5 and 0.005 *μ*M Cmp58, respectively, when neutrophils were challenged with acetate. Whereas neutrophils exposed to 0.5*μ*M Cmp58 during the whole procedure were fully responsive to acetate, no neutrophil response was induced by acetate when the concentration of the allosteric modulator was rapidly reduced (from 0.5 to 0.005*μ*M) through the dilution procedure (Fig 5C inset), and the priming effect disappeared already after 30 seconds. Our data thus show that the Cmp58 effect is reversible and, more importantly, that one of the mechanisms suggested to be involved in neutrophil priming could be excluded. In many studies a direct link has been described, between priming and recruitment to the cell surface, of receptors stored in easily mobilized secretory organelles (38, 39), and such a mobilization is irreversible (40).

### No inhibition of down-stream signaling of FFA2R is induced/mediated by the selective G*α*q inhibitor YM-254890

It has been suggested/shown that FFA2R can couple to both G*α*i and G*α*q for down-stream signaling (17, 41, 42) suggesting that a possible mechanism for the priming effect of the allosteric modulation of Cmp58 on the FFA2R response could be at the level of the signaling G-protein. To elucidate this possibility, we applied the selective G*α*q inhibitor YM-254890 which inhibits the down-stream signaling of the neutrophil PAFR whereas no effect is obtained on the response induced by agonist occupied FPRs (Fig 6A and (43)). The G*α*q inhibitor YM-254890 had no major effect on Cmp1/FFA2R mediated NADPH-oxidase activity, and the inhibitory profile was the same in Cmp58 primed neutrophils (Fig 6B and C). It has been well established that neutrophil chemotactic GPCRs constitute a receptor class that is coupled to a pertussis toxin-sensitive G protein (44), and we show that also the FFA2R response was sensitive to pertussis toxin treatment. The cells treated with pertussis toxin were thus non-responding to acetate/Cmp1 combined with Cmp58 but fully responsive to PMA (a ROS inducer that signals independent of a G-protein; Fig 6D).

**Figure 6.**
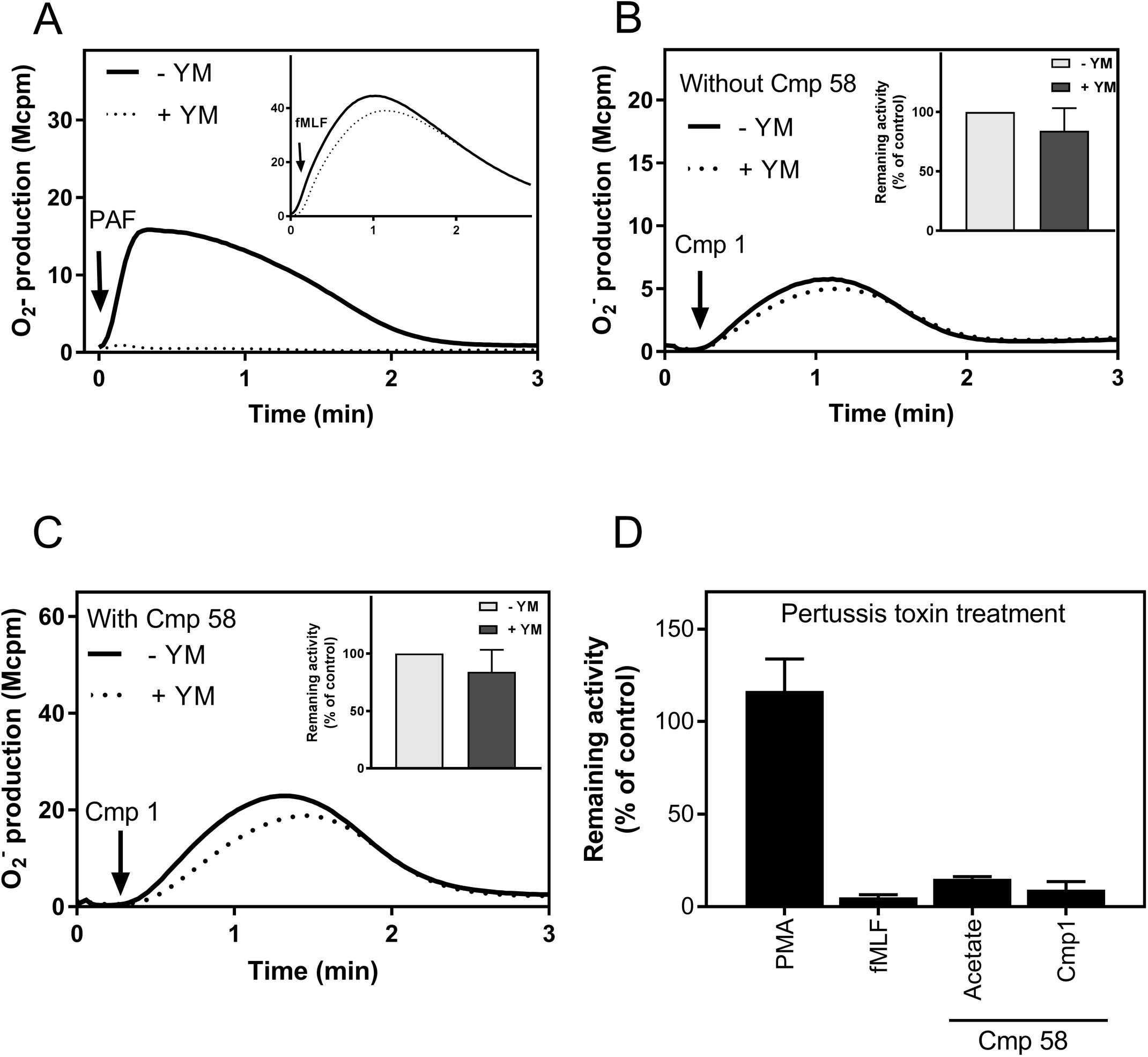
The FFA2R induced activation of the neutrophil NADPH-oxidase does not engage Gαq signaling but is inhibited by pertussis toxin. A. To show the receptor selectivity of the Gαq inhibitor YM-254890, human neutrophils were incubated for 5 min (at 37^°^C) with (dotted lines) or without (solid lines) the the G*α*q inhibitor YM-254890 (200 nM), the cells were then activated by the addition of PAF (100 nM) or fMLF (100 nM; inset) and the release of O_2_^−^ was measured over time. Time point for addition of agonist shown by the arrows. One representative experiment is shown (see also (43)). Abscissa, time of study (min); ordinate, O_2_^−^ production given in arbitrary units (Mcpm). B. Human neutrophils were incubated for 5 min at 37^0^C with or without the Gαq inhibitor YM-254890 (200 nM) and the cells were then activated by the addition of Cmp1(1 *μ*M) at the time point shown by the arrow and the release of O_2_^−^ was measured over time. One representative experiment out of > 3 is shown. Abscissa, time of study (min); ordinate, O_2_^−^ production given in arbitrary units (Mcpm; 10^6^ counts/min). Inset: The inhibitory effect of YM-254890 (200nM) on the Cmp1 induced of O_2_^−^ production, presented as percent of the response (peak value) in the absence of the inhibitor (mean ± SD; n = 3). C. Human neutrophils were incubated for 5 min (at 37^0^C) with Cmp58 together with (dotted line) or as control without (solid line), the Gαq inhibitor YM-254890 (200 nM). The cells were then activated by the addition of the FFA2R agonist Cmp1(1 *μ*M) at the time point shown by the arrow and the release of O_2_^−^ was measured over time. One representative experiment out of three is shown. Abscissa, time of study (min); ordinate, O_2_^−^ production given in arbitrary units (Mcpm). Inset: The inhibitory effect of YM-254890 (200nM) given in percent of the control the Cmp1 responses induced in the absence of the inhibitor. The peaks of O_2_^−^ produced were compared (mean ± SD; n = 3). D. Human neutrophils were incubated without or with Pertussis toxin (PTX, 500 ng/ml) for 150 min at 37°C, the cells were then pre-treated with Cmp58 (1 *μ*M) for 5 minutes before stimulation with Cmp1 (1 *μ*M) or acetate (1 mM). For comparison, PTX treated and untreated neutrophils were activated by fMLF (100 nM) or PMA (50 nM; a non-G-protein dependent stimulus). The peak superoxide activities in the presence and absence of PTX were compared, and the bar graph shows remaining activity following PTX-treatment in percent of the response induced by the agonists in non-treated neutrophils (n = 3; mean±SD).

The fairly limited effect of the G*α*q inhibitor was seen also when looking at the rise in [Ca^2+^]_i_ induced by acetate in the absence and presence of Cmp58 (Fig 7) evidently demonstrating that G*α*q does not play a role in the activity induced in neutrophils by FFA2R agonists.

**Figure 7.**
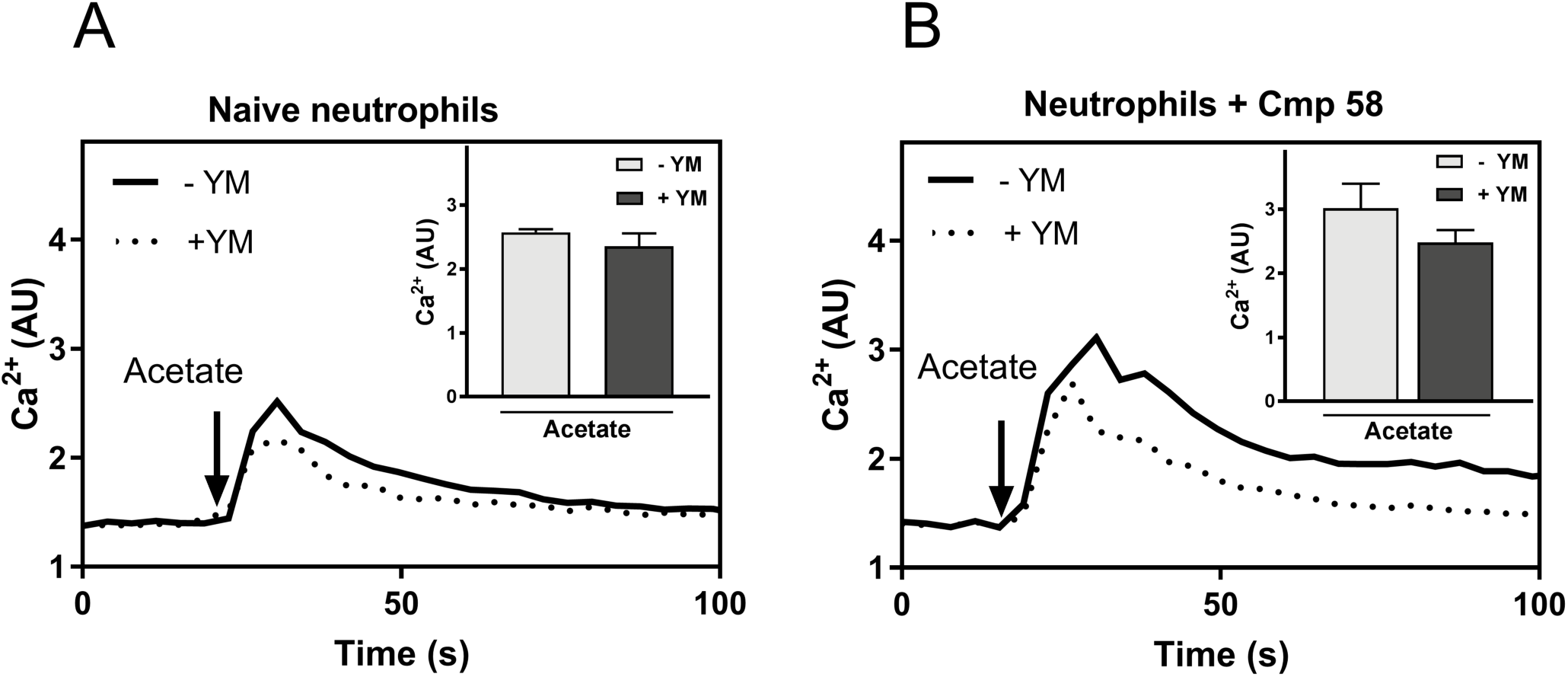
The acetate induced rise in [Ca^2+^]_i_ is not affected by Gαq inhibitor YM-254890 independent of the presence or absence of the allosteric modulator Cmp58. A. Human neutrophils labeled with Fura-2 were used to measure Ca^2+^ transients. The cells were pre-incubated at 37°C for 10 min with (dotted line) or without (solid line) the Gαq inhibitor YM-254890 (200 nM). The neutrophils were then stimulated with acetate (2.5mM; as indicated by the arrow) and the Ca^2+^ transient was measured over time. One representative experiment out of three is shown. Abscissa, time of study; ordinate, increase in intracellular [Ca^2+^]_i_ given by the ratio between Fura-2 fluorescence at 340 and 380 nm. Inset: The bar graph shows the peak values of acetate induced intracellular [Ca^2+^]_i_ in the absence (light grey bar) or presence of the YM-254890 (200 nM dark grey bar; mean±SD; n = 3). B. Neutrophils labeled with Fura-2 were used to measure Ca^2+^ transients. The cells were pre-incubated at 37°C for 10 min with Cmp58 (1 *μ*M) combined with (dotted line) or without (solid line) the Gαq inhibitor YM-254890 (200 nM). The neutrophils were then stimulated with acetate (0.25mM, as indicated by the arrow) and the Ca^2+^ transient was measured over time. One representative experiment out of three is shown. Abscissa, time of study; ordinate, increase in intracellular [Ca^2+^]_i_ given by the ratio between Fura-2 fluorescence at 340 and 380 nm. Inset: The bar graph shows the peak values of cmp58+acetate induced intracellular [Ca^2+^]_i_ induced in the absence (light grey bar) or presence of the inhibitor YM-254890 (200nM, dark grey bar; mean ± SD; n = 3).

### TNF-*α* further primed the Cmp58/acetate-triggered activation of the neutrophil NADPH-oxidase

Neutrophils treated with the pro-inflammatory cytokine TNF-*α* exposed an increased number of receptors such as complement receptor 3 (CR3; CD11b) i.e., TNF-*α* induces mobilization of receptor storing organelles and an increased exposure of receptors to the cell surface (Fig 8A and (23). No NADPH-oxidase assembly and activation was obtained when neutrophils were challenged with TNF-*α* and this is in agreement with our earlier reported results (23). In addition, no oxidase activity was induced in TNF-*α* treated cells by Cmp58 or acetate when added alone in concentrations up to 1*μ*M and 1 mM, respectively (data not shown). TNF-*α* was, however, an additional priming agent when the two FFA2R ligands were combined (Fig 8B). Previous studies have shown that many neutrophil chemoattractant receptors are stored in secretory organelles of naïve cells (38, 45, 46), suggesting that mobilization of these receptors could be a contributing mechanism in priming. However, no increased surface exposure of FFA2R was induced by TNF-*α* (Fig 8C) indicating that FFA2R is not stored in TNF-*α* mobilized neutrophil granules.

**Figure 8.**
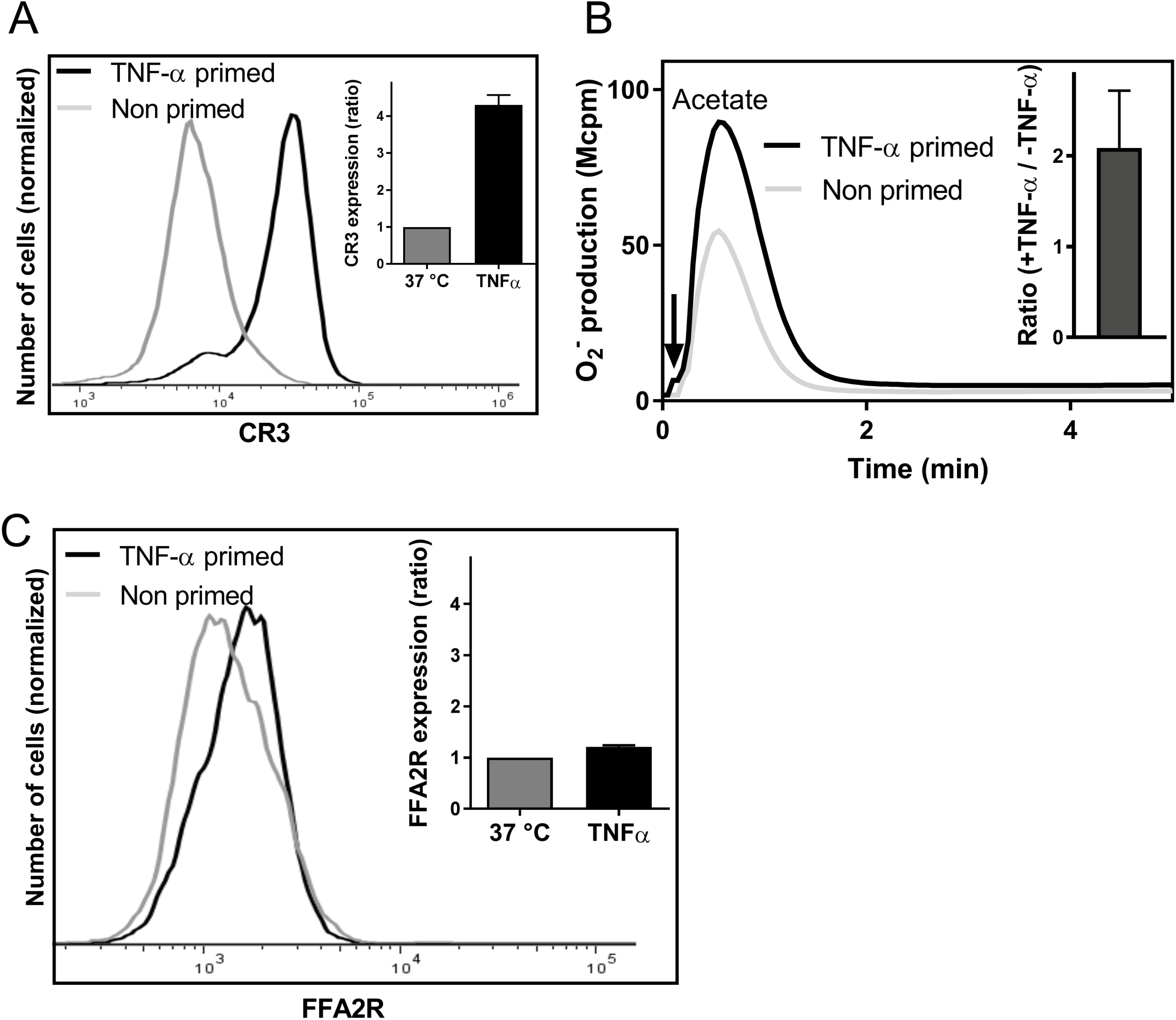
TNF-*α* further primes the neutrophil response induced by acetate in the presence of the allosteric modulator Cmp58. A. The histogram shows the neutrophils cell-surface expression of CR3 after incubation without (light grey) or with TNF-*α* (10 ng/mL) for 20 min at 37°C, as measured by flow cytometry. Inset: The bar graph shows the mean fold increase in fluorescence for primed cells (TNF-*α*, black bar) as compared to naïve (non-primed) cells (grey bars; mean±SD; n=3). B. Neutrophils pre-incubated for 15 min at 37°C without (dotted line) or with (solid line) TNF-*α* (10 ng/ml) were then incubated for another 5 minutes with Cmp58 (1*μ*M) and activated with the FFA2R agonist acetate (1mM). Superoxide production was recorded continuously and one representative experiment out of three is shown. Abscissa, Time of study (min); Ordinate, Superoxide production (peak value in arbitrary units; Mcpm). Inset: The peak NADPH-oxidase values were determined and the bar graph shows the ratio between the value obtained with TNF-*α* primed cells compared to that obtained with naïve (non-TNF-*α*–treated) cells (mean±SD; n=3). C. The histogram shows the neutrophils cell-surface expression of FFA2R after incubation without (light grey) or with TNF-*α* (10 ng/mL) for 20 min at 37°C, as measured by flow cytometry. Inset: The bar graph shows the fold of FFA2R expression for primed cells (TNF-*α*, black bar) as compared to non-primed cells (grey bars; mean±SD; (n=3).

## Discussion

The allosteric FFA2R modulator Cmp58 ((S)-2-(4-chlorophenyl)-3,3-dimethyl-N-(5-phenylthiazol-2-yl)butanamide) primes neutrophils in their response to acetate as well as the other natural FFA2R agonists propionate and butyrate. The molecular mechanisms underlying neutrophil priming has been extensively studied and a large number of different mechanisms have been suggested (38, 47–52). By showing that Cmp58 changes the natural FFA2R agonist acetate from being non-activating to a potent neutrophil activating agonist that triggers an assembly of the superoxide generating NADPH oxidase and lowers the thresh-hold for the FFA2R-induced intracellular Ca^2+^ release, we add allosteric receptor modulation as a novel receptor specific neutrophil priming mechanism. The priming agent Cmp58 (22) has earlier been shown to be an FFA2R modulating agonist that binds to an allosteric site on the receptor, and it has also been demonstrated that binding of Cmp58 co-operatively modulate the activity of natural receptor specific ligands in cells other than neutrophils (22). This type of co-operativity between the allosteric agonist Cmp58 and the orthosteric agonist acetate fits with the neutrophil priming data described in this study, except that no direct neutrophil activation was obtained with the allosteric compound in the concentration range tested. We have earlier shown that FFA2R agonists cross desensitizes each other (23); the Cmp58 induced priming is thus not due to an additive effect of two FFA2R agonists. The suggested mechanism for how allosteric modulators affect receptor function involves a direct interaction of the modulator with a specific allosteric binding site and this interaction secondarily transfers the receptor either i) to a state possessing a lower energy barrier for the conformational change required for a shift from a resting non-signaling receptor state to an activated signaling state and by that an increased signaling activity downstream of the agonist occupied receptor, ii) to a state in which the orthosteric binding site has a higher affinity in binding to conventional agonists or, iii) through a combination of the first two (53). It is clear from the dilution experiment performed (see Fig 5C) that the dissociation of acetate from its receptor is very rapid, suggesting that it is very hard to perform conventional binding experiments with this receptor agonist (54). It has, however, been shown, using a three-way radio-ligand binding experiment, that the ability of acetate to inhibit receptor-binding of an FFA2R antagonist ([^3^H]GLPG0974) is increased by allosteric FFA2R modulators/agonists (41). Based on these findings it was suggested that an increased binding affinity for orthosteric FFA2R agonists is part of the allosteric effect (41), but irrespectively of the precise modulation mechanism for Cmp58, the outcome of the modulating effect in neutrophils is that acetate is turned into an agonist that activates neutrophils to generate and release superoxide anions.

There has been an urgent need for receptor specific tools to allow studies of FFA2R in neutrophils, and among the ligands identified (29), the agonist/antagonist pair Cmp1 and CATPB, fulfilling the requirements of being both potent and receptor selective, were recently used by us to disclose the basic functional characteristics of FFA2R in phagocytic cells (23). Briefly, we showed that Cmp1 triggered an increase in the cytosolic concentration of Ca^2+^ and following this, neutrophils were desensitized not only to Cmp1 but also to the naturally occurring FFA2R agonist acetate, and the antagonist CATPB selectively inhibited the responses. Moreover, the functional and regulatory differences between FFA2Rs and another group of pattern recognition receptors (i.e., the FPRs) imply different roles of these receptors in the orchestration of inflammation and confirm the usefulness of selective FFA2R agonists and antagonists as tools for the exploration of the precise functions of FFA2R. Activation of neutrophils with the FFA2R agonist Cmp1 induced an assembly of the superoxide generating NADPH-oxidase but no chemotaxis, and despite an induction of a rise in intracellular Ca^2+^ by Cmp1, this response was not accompanied by any mobilization of secretory vesicle proteins or cleavage of L-selectin from the cell surface, the two most sensitive markers for neutrophil activation. We now introduce an allosteric modulator as a new receptor specific tool that can be used to study FFA2R in neutrophils. A new concept in the regulation of GPCR signaling was introduced when allosteric ligands were identified and shown to affect (positive or negative) receptor functions when triggered by conventional ligands that interact with the orthosteric binding site exposed by the targeted receptor (53, 55). This concept also adds complexity to the pharmacology and signal transduction scheme downstream an activated receptor, and many different models of how to categorize and analyze this type of allosterism has been described (28, 30). It is clear from the data presented that the FFA2R selective positive allosteric modulator used by us (Cmp58) as well as other modulators such as 4-CMTB (56) and AZ1729 (12) affects the response induced by FFA2R specific orthosteric agonists. The 4-CMTB and AZ1729 compounds have been shown to be both agonists and modulators when interacting with mouse and human neutrophils, respectively (41, 56). The lack of direct neutrophil activating effects of Cmp58/4-CMTB in our systems might be due to species differences, shown to be of importance both for ligand recognition and for signaling of other GPCRs (6, 57) or to differences between the modulating compounds and/or the read-out systems used. Although 4-CMTB is an allosteric modulator it has been shown to have unique properties (24) that could possibly explain why the positive allosteric effect mediated by 4-CMTB on the neutrophil response induced by Cmp1 (Fig 5D) is not evident when determined in other cells or read-out systems (20). The maximal modulating effect of Cmp58 was reached at a concentration of 1*μ*M, suggesting that the allosteric binding sites are fully occupied at this concentration. The lack of direct neutrophil activating effects of Cmp58 when added alone even at a concentration of 5*μ*M, suggests that it should be classified as an allosteric modulator (rather than an allosteric agonist) in relation to FFA2R when the receptor is expressed in neutrophils.

The response induced in neutrophils by Cmp58/acetate was abolished in pertussis toxin treated neutrophils, with the logical conclusion that FFA2R signals through a Gαi containing G-protein. We have, however, recently shown that also the neutrophil response mediated by the agonist occupied Gq-coupled receptors for platelet activating factor is inhibited by pertussis toxin, showing that pertussis toxin experiments are not always reliable (43). It has been suggested/shown that FFA2R can couple not only to G*α*i but also to G*α*q for down-stream signaling (17, 41, 42), but the Cmp58/acetate response was insensitive to the Gq inhibitor YM-254890. Taken together, these data suggest that the signaling cascade triggered by FFA2R and leading to a rise in intracellular Ca^2+^ and activation of the NADPH-oxidase in neutrophils do not involve G*α*q, but rather is initiated by a Gαi containing G-protein.

Irrespectively of the precise molecular mechanism for the Cmp58 mediated modulation of FFA2R, the functional outcome is that the neutrophil response induced by conventional receptor agonists is primed, and this was evident also in TNF-*α* treated cells. It is clear from earlier studies that a prominent function of TNF-*α* is the capacity to in itself prime neutrophils in their response to other stimuli, and we have earlier suggested that mobilization of receptor storage organelles that is induced by the cytokine is an important mechanism in neutrophil priming. Our data showing a lack of FFA2R mobilization in TNF-*α* primed neutrophils do, however, not support this suggestion. It is clear that the receptor storing organelles mobilized in TNF-*α* primed neutrophils contain also the flavocytochrome b_558_ (58), the membrane component of the superoxide generating NADPH-oxidase, and mobilization of this protein may well be part of the TNF-*α* priming process (59). Since augmentation mediated by Cmp58 is reversible, whereas mobilization of neutrophil secretory organelles achieved through a fusion of granule/vesicle membranes with the plasma membrane is a non-reversible process that stably mobilizes proteins from granule/vesicle membranes to the plasma membrane (40), receptor mobilization is not a process directly involved in the Cmp58 induced priming. Regardless of the precise mechanisms for TNF-*α* priming it is clear that the allosteric modulator adds to the TNF-*α* effect in making acetate an even more potent NADPH-oxidase activating agonist. The type of allosteric modulation presented in this study, is a novel avenue for potential regulating effects of G-protein coupled receptors expressed in neutrophils. They may offer greater receptor selectivity by binding to non-conserved regions of the receptor among a family of related receptors that may offer other potential advantages such as differential effects on tachyphylaxis and efficacy. Positive allosteric modulators/agonists thus offer an attractive therapeutic approach for the regulation of GPCRs (30). It is clear from the data presented that the neutrophil response is affected by Cmp58 when triggered by sub-optimal concentration of an FPR1 agonist. There are several possible mechanisms to this effect, including a receptor dimerization that has been described to modulate fatty acid sensing by FFA2R/FFA3R heterodimers when expressed in monocytes/macrophages (60). The fact that no neutrophil response is induced by the FFA3R agonist AR420626 support an earlier finding, namely that this receptor is not expressed in neutrophils (17), but this does not exclude receptor cross talk between FFA2R and other receptors expressed in these cells. The precise mechanism by which Cmp58 affects the neutrophil response to non-activating concentrations of the FPR1 agonist, thus, needs to be studied in more detail.

We show that Cmp58, an allosteric modulator for FFA2R, triggers signals generated in neutrophils by the combined effect of the modulator and the natural agonist acetate determined as release of superoxide anions. This response is even further increased in TNF-*α* primed neutrophils. The fact that short chain fatty acids are produced by gut microbes during anaerobic metabolism, has suggested that receptors for such molecules expressed on innate immune cells are of prime importance for regulating inflammatory reactions in the gut (12). However, the fact that tissue-recruited neutrophils are non-responding to the FFA2R agonist (23), suggests a role for the receptor also in other organs implying that the role of FFA2Rs is complex and needs further investigation. The phenylacetamides discovered as a novel class of FFA2R allosteric modulators, positively co-operate with natural FFA2R agonists and prime neutrophils in their response to these agonists, may serve as good tools for further unraveling the physiological functions of the FFA2R and its involvement in various diseases. In addition, and even though the majority of the allosteric modulators described so far have been identified through high-throughput screening processes using different libraries of small molecules, the fact that the natural ligands for FFA2R are free fatty acids generated by the fermenting microbiota in the gut, unavoidably raises the question whether endogenous or other naturally occurring microbial metabolites can fulfill an allosteric function and physiologically change the functional repertoire of FFA2R and other GPCRs.

## Abbreviations

BSA = bovine serum albumin; CATB = the FFA2R antagonist 3-benzyl-4-(cyclopropyl-(4-(2,5-dichlorophenyl) thiazol-2-yl)amino)-4-oxobutanoic acid]; CL = chemiluminescence; Cmp1 = the FFA2R agonist 3-benzyl-4-(cyclopropyl-(4-(2,5-dichlorophenyl) thiazol-2-yl)amino)-4-oxobutanoic acid]; Cmp58 = the allosteric modulator (S)-2-(4-chlorophenyl)-3,3-dimethyl-N-(5-phenylthiazol-2-yl)butanamide; CR3 = complement receptor 3; DAMP = danger associated molecular pattern; FFA2R = free fatty acid receptor 2; FFA3R = free fatty acid receptor 3; fMet = formyl-methionyl; fMLF = formyl-Met-Ley-Phe; FPR = formyl peptide receptor; GPCR = G-protein coupled receptor; GPCR41 = free fatty acid receptor 3/FFA3R; GPR43 = free fatty acid receptor 2/FFA2R; GPR84 = G-protein copled receptor 84; HRP = horse radish peroxidase; IP_3_ = inositol-1,4,5-tris-phosphate; KRG = Krebs-Ringer Glucose phosphate buffer; MCPM = mega-counts per minute; O_2_^−^ = superoxide anion; PAMP = pattern associated molecular pattern; PLC; phospholipase C; PIP_2_ = phosphatidyl-inositol-4,5-bis-phosphate; PPR = pattern recognition receptors; SCFA = short chain fatty acid; TNF-*α* = Tumor necrosis-*α*; WKYMVM = Trp-Lys-Tyr-Met-Val-L-Met; YM-254890 = a G-protein*α*q inhibitor; 4-CMTB = the allosteric agonist/modulator 4-Chloro-α-(1-methylethyl)-N-2-thiazolyl-benzeneacetamide

## Acknowledgment

The work was supported by the Swedish Medical Research Council (CD, 005601; HF, 02448) the King Gustaf V 80-Year Foundation (CD FAI 2014-0011), the Clas Groschinsky Foundation (HF, MI562), the Wilhelm and Martina Lundgrens Scientific Foundation (MS 2017-1925), Åke Wibergs Foundation (HF, M15-005), and the Swedish state under the ALF-agreement (CD, ALFGBG 72510).

Valuable suggestions provided by the members (past and present) of The Phagocyte Research Group at the Sahlgrenska Academy, University of Gothenburg, and the experimental work performed by Linda Bergqvist are gratefully acknowledged.

## Authorship contributions

JM and AH designed and performed research and analyzed data. MS, MG and LB performed research and analyzed data. HF and CD planned and supervised the research, analyzed data, and wrote the manuscript that was edited by all authors.

